# Cleared tissue dual-view oblique plane microscopy

**DOI:** 10.1101/2025.05.12.653537

**Authors:** Liuba Dvinskikh, Hugh Sparks, Darren Ennis, Raffaele Sarnataro, David Carreno, David T. Riglar, Paula Cunnea, Iain A. McNeish, Chris Dunsby

## Abstract

We present a dual-view oblique plane microscope (dOPM) for imaging thick optically cleared tissue samples using a silicone immersion primary objective. The custom-designed remote refocusing relay utilizes stock optics to achieve remote refocusing in refractive index-matched samples. The spatial resolution of the system was characterized using a series of fluorescent bead phantoms with refractive indices ranging from 1.4-1.5, with the point spread function full width at half maximum measuring ∼0.5 µm laterally and ∼1 µm axially for a refractive index-matched bead sample, with minimal degradation over a >250 µm axial range. We characterize how the remote refocusing performance at sample refractive indices up to n = 1.5 can be partially compensated for using adjustment of the correction collar on the primary objective. We apply the system to imaging a range of biological samples with varied refractive indices. Combined with tiled acquisition, image stitching and multi-view image fusion, the microscope enables multicolour imaging of mm-wide and ∼250 µm thick optically cleared mouse ovarian cancer and colon tissue samples with subcellular resolution. We also image a whole *Drosophila melanogaster* fruit fly brain. The system offers a platform for fast and high-resolution, multicolour volumetric imaging across spatial scales, integrated on a commercially available microscope frame.

## 1. Introduction

The comprehensive study of biological structure and function in context of the complex microenvironment has driven the development of imaging techniques capable of visualizing larger multicellular three-dimensional (3D) samples such as tissues and organoids. The optical sectioning of light-sheet fluorescence microscopy (LSFM) offers a non-destructive alternative to physically-sectioning histopathology-based techniques for visualizing the 3D architecture of these samples.

A key challenge in imaging larger structures is the limited optical imaging depth, with scattering as the main mechanism of signal attenuation. Scattering arises due to a non-uniform refractive index (RI) distribution, with higher-RI biomolecules such as lipids and proteins surrounded by lower-RI aqueous solution. Optical clearing aims to decrease this RI mismatch, reducing sample scattering and improving image resolution and contrast, with the final RI of cleared tissue typically ranging between 1.38-1.56 [1]. Today, a wide range of clearing protocols exist, including solvent based, immersion, hyperhydration and hydrogel embedding methodologies. In simple immersion, the sample is gradually cleared by placing the tissue in an aqueous solution of a high-RI molecule. A low-cost non-toxic option is 2,2’-thiodiethanol (TDE), allowing linear RI-tuning over the 1.33-1.52 range by mixing with water at various ratios [2], [3], [4]. However, in addition to RI variation within the sample, RI mismatch between the sample and immersion medium results in focal shift, axial distortion and loss of image contrast and resolution due to depth-dependent spherical aberrations [5], [6], hence there is a need to ensure compatibility between the imaging optics and clearing protocol.

A further key consideration in the imaging of thicker tissues is sample mounting. In conventional dual-objective LSFM configurations, one objective is used to create a thin sheet of light to excite fluorescence from a single slice of the sample, while a second, orthogonally oriented objective collects that fluorescence. The two closely positioned objectives introduce steric hindrance constraints on sample mounting. An open-top light-sheet microscope (OTLS) [7] enables the specimen to be mounted on a glass or plastic plate with illumination and detection through the plate. OTLS has been used for example to demonstrate non-destructive pathology of large clinical specimens using a fluorescent analogue to haematoxylin and eosin (H&E) staining [8]. However, the angled objective orientation relative to the sample-mounting plate reduces the usable working distance of the objective and limits the numerical aperture (NA).

Oblique plane microscopy (OPM) [9] achieves light-sheet excitation of a tilted plane in the sample, and orthogonal detection of the emitted fluorescence through a single high-NA primary objective lens, enabling conventional sample mounting using coverslips or multiwell plates. In OPM, two back-to-back microscope relays (with primary objective O1, tube lenses TL1 and TL2, and secondary objective O2) are used to generate an isotropically magnified image through remote refocusing [10], [11], with a third, tilted, microscope relay (with tertiary objective O3 and tube lens TL3) imaging the illuminated sample plane onto the camera. Aberration-free remote refocusing is achieved when the pupil of O1 is imaged onto the pupil of O2 such that the rays from points in object space converge stigmatically to a point in image space and that the ray angles in object and image space are equal. This conservation of conjugate ray angles requires that the lateral and axial magnification of the remote refocusing relay is equal to the ratio of the RIs of sample and image space.

Previous characterization of remote-refocusing relays have shown that even small deviations from the target magnification and axial misalignment of O1, O2, TL1 or TL2 result in a significant decrease in the diffraction-limited remote-refocusing range [12], [13]. Additionally, it has been shown that away from the focal plane, residual spherical aberrations arise even in a well-aligned remote-refocusing relay, and the diffraction-limited range can be enhanced through adjustment of the correction collar (CC) of O1 [14].

An oblique plane in the sample can also be imaged using a folded remote-refocusing arrangement with a tilted mirror in the focal plane of the second objective O2, allowing the second objective to be used twice, functioning as both O2 and O3 [15], [16]. Folded remote refocusing is used in dual-view oblique plane microscopy (dOPM) [17], where the translation of a pair of tilted mirrors in refocusing space enables the imaging of two orthogonal views of the sample, with the dual-view configuration reducing shadow artefacts and enabling a more isotropic resolution.

A key challenge in using OPM for imaging of larger samples is that the lower-magnification and larger field of view (FOV) O1 required to visualize large structures tend have a lower NA, and re-imaging the inclined plane from a low or moderate NA OPM system involves higher loss of the light-cone with higher tilt of the third microscope relay. This can be overcome using a high-NA zero working distance tertiary objective [18], which can also be combined with mirror-based beam rotation for multi-view imaging [19], however the optical train is longer and involves additional pupil relays. Using a non-orthogonal dual-objective light-sheet configuration [20] with a vertical detection objective, and non-orthogonal illumination provided by a separate objective, reduces aberrations in the detection path compared to a orthogonal objective OTLS approach, and allowing the use of a moderate-NA primary lens, but with a shallow excitation angle. A similar excitation approach is used in direct-view OPM [21], where the collection efficiency of a low-NA OPM is further increased by positioning a camera directly in the remote image space after O2.

In this work, we present the design and use of a dual-view OPM system employing a silicone immersion primary objective (O1) and apply it to imaging optically cleared tissues. We provide a detailed measurement of the system performance using experimental PSF measurements in RI-tuned bead phantom samples over the range n = 1.4-1.5. We also provide a detailed characterisation of the effect of using the primary objective correction collar when imaging samples with different refractive indices. We demonstrate proof-of-concept fast, high-resolution, multicolour biological imaging of few hundred µm thick samples with RI values in the aforementioned range.

## 2. Methods

### 2.1. Optical system configuration

The system design has been adapted from the dual-view oblique plane microscope presented by Sparks et al. [17], with a modified folded remote refocus relay operating with a silicone-immersion O1 (Nikon CFI Plan Apo Lambda S40XC Sil, 1.25 NA) (**Figure *1***). The sample is mounted on a motorized XY-stage (SCANplus IM 130 × 85, Marzhauser). The first microscope consists of the primary objective and 200 mm Nikon tube lens (TL1) housed in a Nikon ECLIPSE Ti2-E frame. A widefield epifluorescence/brightfield optical path set up through the right-hand camera (ORCA-Fusion, C14440-20UP, Hamamatsu) port was used for alignment and sample navigation. The intermediate image plane of the remote refocus relay is located just outside of the left-hand camera port, and is re-imaged by a second microscope relay consisting of a tube lens (TL2) formed by two air spaced doublets (see **section 2.1.1.1** for details), paired with an Olympus UPLXAPO 20x 0.8 NA air objective (O2/3) with a #1.5 glass coverslip glued to the front lens (mounted as a service by ASI). Two dielectric-coated mirrored prisms, angled at ±22.5 degrees relative to the optical axis of O2/3 are placed in the focal plane of O2/3. The mirrored prisms are held in a custom common mount on a motorized linear stage (PIMag, V-522.1AA, Physik Instrumente) on top of an orthogonally oriented micrometer translation stage (UMR5.16, Newport). The same objective O2/3, paired with a third tube lens TL3 (TTL200-A) is used to image the mirror surface in remote image space onto a second sCMOS camera (ORCA-Fusion), through a motorized emission filter wheel (EM, FF01-525/45-25, FF01-600/52-25, FF01-698/70-25 emission filters, Brightline, Semrock, paired with 488 nm, 561 nm and 642 nm laser excitation lines respectively). A polarising beam splitter cube (PBS, CCM1 PBS251/M, Thorlabs) and a quarter-wave plate (QWP2, AQWP10M-580, Thorlabs) and periscope mirror M6, positioned between TL2 and O2 are used to couple the detected light out of the remote refocusing relay, with QWP2 set to maximize transmission of green-filtered detection light from the sample to the camera through the PBS.

**Figure 1.**
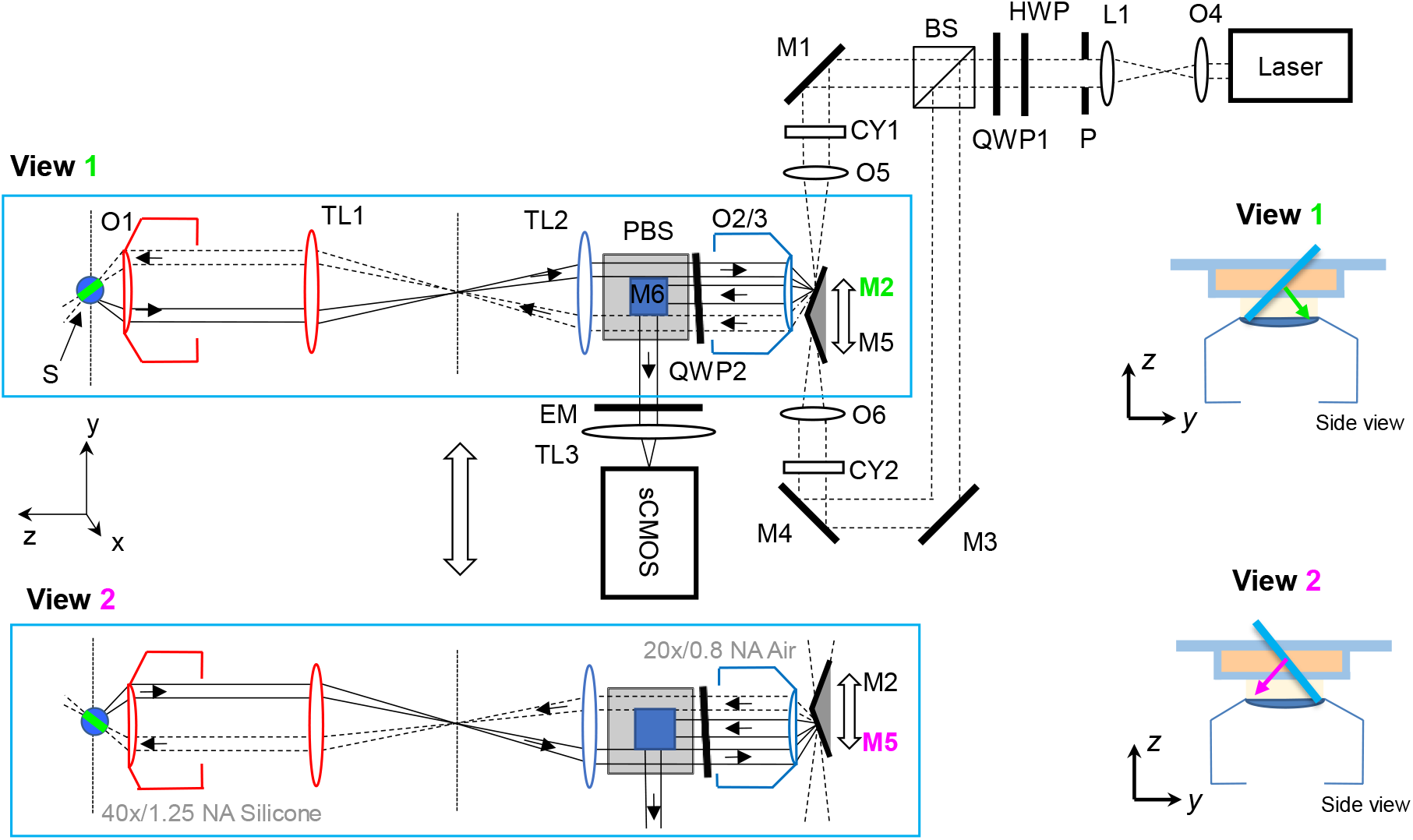
Optical system configuration. S, sample; O, microscope objective; TL, tube lens; PBS, polarising beam splitter; BS, non-polarising beam splitter; HWP, half-wave plate; QWP, quarter-wave plate; M, mirror; CY, cylindrical lens; P, pinhole; L, lens; EM, emission filter. Top part of figure, optical configuration for View 1, bottom part of figure, optical configuration for View 2. Insets on right, closeup schematic of orientation of illumination light sheet in sample for both views.

Excitation light provided by a multi-wavelength CW laser engine (Omicron LightHUB) is expanded by objective O4 (Olympus UPlanFL N 10x/0.3NA) and lens L1 (AC508-200-A-ML, Thorlabs), passed through a 4.5 mm diameter pinhole, and split into two paths by a 50/50 beam splitter (BS, CCM1-BS013, Thorlabs). The excitation light in the two paths is folded by mirrors M1, M3, M4 and focused down to a line by 150 mm focal length cylindrical lenses CY1&2 (LJ1629RM-A, Thorlabs) onto the back-focal plane of air objectives O5&6 (Olympus UPLFLN 4X) to create a light-sheet in the focal plane of O2/3. The tilted mirrors M2&5 couple the light-sheet into O2/3 at ±45 degrees to its optical axis, through the remote refocusing relay to the sample, and their translation orthogonal to the optical axis is used to swap between the two views and synchronously sweep the light-sheet and detection plane through remote-space, scanning a volume through the sample. A quarter-wave plate QWP1 is used to generate a circularly polarized beam, which is converted back to linear by QWP2, with the axis of polarization set by the half-wave plate (HWP, AHWP10M-580, Thorlabs) earlier in the path to maximize transmission through the PBS to the sample.

#### 2.1.1. Aberration-free remote refocusing detection path

##### 2.1.1.1. Design of custom tube lens TL2 from stock optics

In order to achieve the desired TL2 focal length for correct magnification from sample to remote refocus space, the second tube lens was constructed using two air-spaced stock achromatic doublets (**Figure 2**), selected using the Doublet Selector automatic lens design software developed by Hong and Dunsby [22]. As is common for commercial microscope frames, O1 and TL1 are not telecentric and hence imaging the pupil of O1 onto the pupil of O2 requires non-telecentric separation between TL2 and O2. The separation *a* (see **Figure *2***) between the back focal plane (BFP) of O1 to the first principal plane (PP1) of TL1 was determined by removing O1, introducing an iris where the back focal plane of O1 is located and measuring the displacement of the image of the iris formed by TL1 from the intermediate image plane (IIP in **Figure *2***). The required distance *b* between the PP1 of TL2 and the O2 BFP was calculated out using the thin lens equation and used in the subsequent optimisation, see below.

**Figure 2.**
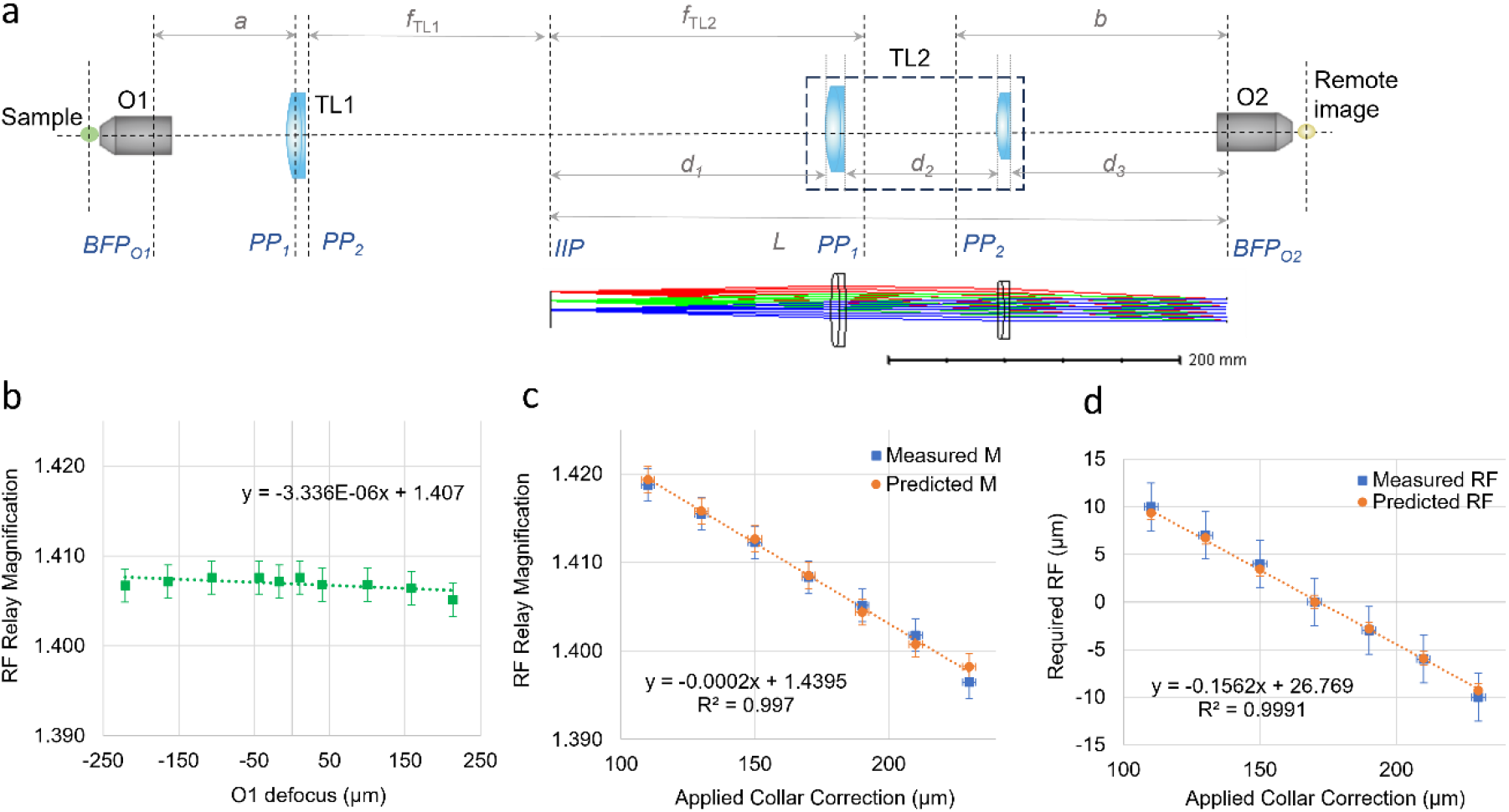
(a) Layout of the remote refocusing relay, consisting of silicone immersion objective O1, tube lens TL1, custom tube lens TL2 composed of two air spaced achromatic doublets, and air objective O2. The distances d_1-3_ indicate the separations between the intermediate image plane (IIP) and the first lens surface, the separation between the two adjacent lens surfaces, and the distance between the final TL2 surface and the back focal plane of O2. The first and second principal planes of TL1 and TL2 are indicated by PP_1_ and PP_2._ The inset below shows the Zemax ray propagation model for TL2. (b) Change in lateral magnification of the remote refocusing relay with axial translation of O1, with minimal variation (gradient <10^-5^ µm^-1^) over >400 µm refocus range. (c) Near-linear change in the measured magnification (blue squares) of the remote refocusing relay with varied CC position, compared to the expected change based on the CC position-dependent focal length of O1 only (orange circles). (d) Comparison of the predicted axial translation of the remote refocusing mirror (orange circles) with the actual measured translation required (blue squares) to refocus on the sample for varied amounts of applied collar correction.

The design parameters corresponding to the required maximum field angle, desired focal length and entrance pupil diameter were calculated using equations 2, 3 and 4 respectively of reference [22], and together with distance *b* used as input parameters for the Doublet Selector software. The selection process was repeated for design parameters corresponding to 5 different magnifications ranging 1.4-1.5, identifying suitable doublet combinations from stock optics. The selection process was performed with the higher curvature surfaces facing in the same direction for both lenses, as this had been found to give the best performance on average across various lens combinations [22]. The final lens selection was made considering the lens combinations with the optimal performance across the magnification range and stock availability, with the selected lens combination consisting of two achromatic doublets with respective focal lengths of 451.5 mm (DLB-500-450PM, Optosigma) and 450.2 mm (L-L-AOC241, Ross Optical), mounted inside an adjustable lens tube (SM2V15, Thorlabs). The approximate lens surface separations were identified using the Zemax model and used as a starting point for the alignment.

#### 2.1.1.2. Alignment and magnification measurement

The optical axis for the second microscope relay was defined using a collimated light beam through the objective turret, with O1 replaced with a pinhole positioned at its BFP. The distance between TL1 and TL2 was set such that the two tube lenses form a 4f system, with the separation adjusted using a collimated alignment laser coupled into the objective turret and a shear plate. Initially, the secondary objective O2 was positioned where the relay composed of TL1 and TL2 form an image of a pinhole placed in the BFP of O1. A flat mirror was mounted at the focal plane of O2/3 normal to the optical axis of O2/3, and the reflected beam was coupled out of the refocusing relay via the PBS and periscope mirror M6 and focused onto the camera by TL3. The quarter waveplate (QWP) is adjusted for maximum transmission of green wavelengths during double pass through the PBS.

With the CC of O1 set to the nominal value (0.17), a transmission 1951 USAF resolution test target (R3L1S4N, Thorlabs) is mounted at the focus of O1 with a #1.5 glass coverslip. The axial position of the plane mirror at O2/3 is adjusted such that the image is simultaneously in focus on both the widefield right hand port optical path and the left-hand port remote refocusing path, and simultaneously in focus with the mirror surface. The lateral position of TL2 was set such that the image of the test target does not translate laterally with remote refocus. The separation of the doublets making up TL2 was adjusted until the desired a remote-refocus (O1, TL1, TL2, O2) magnification of 1.406 is achieved, after which the position of O2 along the optical axis is iteratively adjusted to minimise the variation of lateral magnification with remote refocus. A gradient of <10^-5^ µm^-1^ for the magnification variation measured over >400 µm refocus range was achieved **(Figure 2b)**. The magnification was determined by measuring the lateral and axial extent of the 3-line features in Group 4 Element 6 of the test target, with a final value of lateral magnification average of M_RF_ = 1.407±0.002. The magnification of the third microscope relay (M_3_ = 22.57) was measured by laterally translating the plane mirror mounted on the PIMag actuator and measuring the position of imperfections on the mirror, giving an overall magnification of M = 31.75, and an effective pixel size of 0.205 µm in sample space.

The adjustment of the CC of O1 over the full range changes the magnification of the first microscope (O1 and TL1) linearly by ∼1.5% between the two extremes (±0.75% in each direction) **(Fig S1**) and introduces a 26 µm focal shift (±13 µm in each direction). Adjustment of the correction collar across the full range enabled tuning of the magnification of the remote refocusing relay between 1.396-1.419 at the zero remote refocus position, in close agreement with the predicted values from the estimated change in the corresponding widefield magnification **(Figure 2c)**. For varied CC positions, the measured axial translation of the remote refocusing mirror required to focus on the mounted sample also closely agreed with the predicted refocus calculated based on the relay magnification and O1 focal plane shift **(Figure 2d)**.

After the remote refocus alignment and magnification measurements were completed, the plane mirror is swapped out for the angled mirrored prisms M2/5, such that an image of a tilted plane through the test chart sample is visible for each view. Finally, the test chart is swapped for a fluorescent slide (Autofluorescent slides, Chroma) and the light-sheet is coupled into the objectives using mirrors M1 & M4.

### 2.2. Image acquisition and processing

Image acquisition was controlled by NIS-elements software (Nikon), with laser triggering via a DAQ box (USB-6343, NI). The image acquisition and reconstruction parameters for the data presented in this paper are summarized in **Table S1**. The raw data was reconstructed using the using the Multi-View Fusion plugin [23], (version 0.10.2) using default settings, implementing the affine transformations described in [17] as well as automatic bead-based co-registration and fusion of the two views. RI-tuned bead phantom volumes (see **Supplementary Note 1** for preparation details) were acquired with identical image acquisition parameters. For tiled FOVs, the individual tiles were acquired with 40-47% overlap and stitched after fusion using the Stitching FIJI Plugin using pairwise and grid collection stitching (snake by rows, right & down) [24] with maximum intensity blending as the fusion method and default settings for all other parameters.

## 3. Results

### 3.1 Remote refocusing performance in various refractive indices

#### 3.1.1. Characterization of the spatial resolution

The spatial resolution was experimentally characterized using refractive-index tuned fluorescent bead phantoms (preparation detailed in **Supplementary Note 1**). Bead phantoms were imaged with 488 nm excitation and the 525/45 nm emission filter, with the O1 focal plane set 150 μm beyond the coverslip into the sample. The acquisition was repeated for 7 different CC values (ranging from 0.11 to 0.23), for bead phantoms with refractive indices tuned to 1.406, 1.45 and 1.5.

The PSF characterization across the FOV over the refocusing range was achieved using a custom PSF quantification pipeline, adapted based on previous versions [25], [26]. The overlapping parallelepiped region of both views in the fused volume was selected for analysis (**Figure 3**, red). Cuboid-shaped volumes around identified bead images were extracted, and 1D Gaussians were fitted to the X, Y, Z and laterally integrated Z* bead image profiles to characterize the lateral and axial PSF FWHM and optical sectioning.

**Figure 3.**
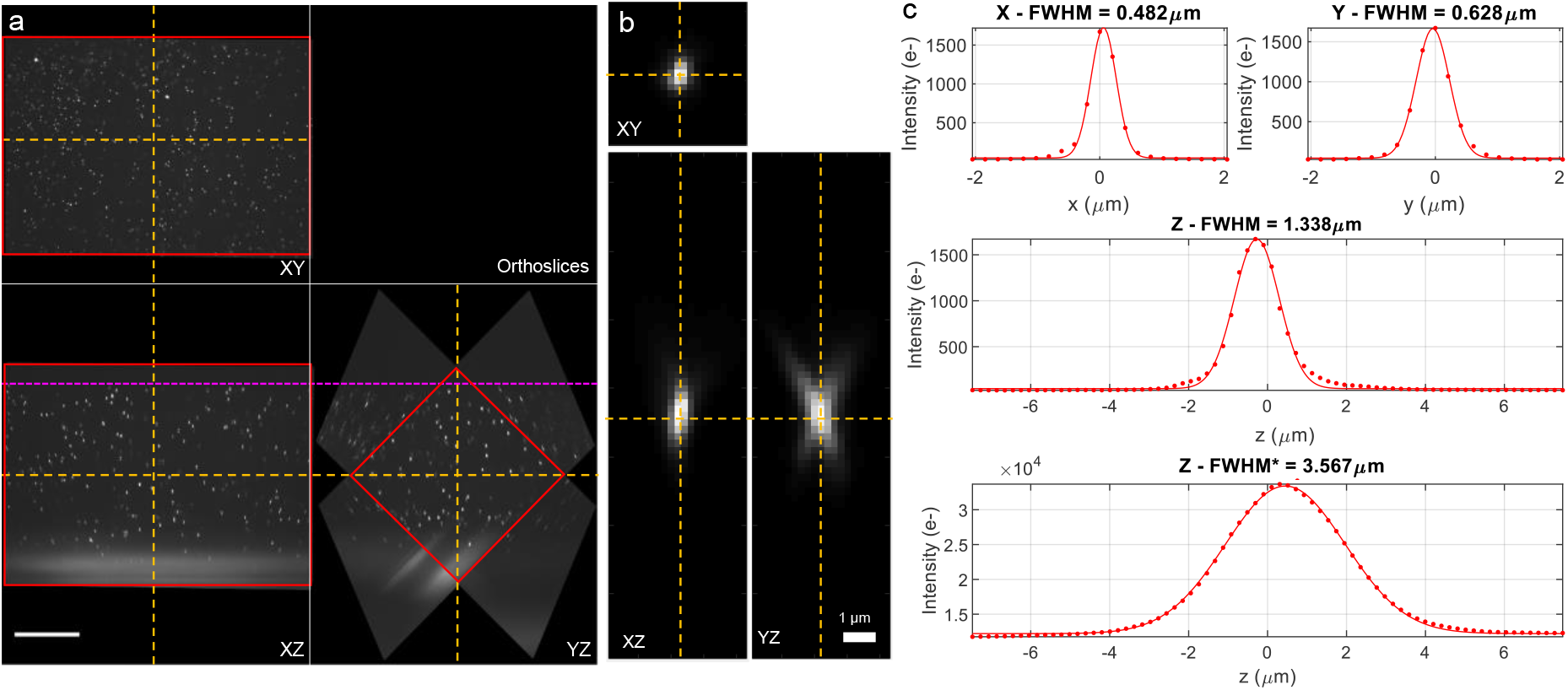
(a) Orthogonal XYZ slices through the central planes (dashed yellow lines) of a fused volume of fluorescent beads in agarose (n = 1.406). The magenta dashed line indicates the coverslip-agarose boundary, with increasing imaging depth in the downwards direction, and the red rectangle indicates the cropped region selected for PSF analysis. Scalebar: 100 µm. (b) Orthogonal slices through the centre of the image of a fluorescent bead. Scalebar: 1 µm. (c) Line profiles and 1D Gaussian fits through the centre of each bead image for the lateral, axial and laterally integrated axial intensity profiles, with the corresponding FWHM.

The resultant values were binned over 3D spatial regions, enabling the generation of three-dimensional heat maps of variation of the lateral and axial resolution as well as optical sectioning across the FOV and with remote refocus.

The experimentally measured spatial resolution with remote refocus for a bead volume with the agarose RI tuned to the design value of 1.406 and CC set to 0.17 is shown in **Figure 4**. The values in **Figure 4a** correspond to the lateral (X and Y), axial (Z) and laterally-integrated axial (Z*) bead image FWHM over the parallelepiped region corresponding to the overlapping volume of both views in the fused data (shown in **Figure 4b**). The lateral and axial resolution remains largely unchanged over a remote refocusing range of ∼250 µm **(Figure 4c)**. The median bead image FWHM over the whole overlapping volume region was measured to be 0.49 ± 0.11 µm and 0.60 ± 0.10 µm in the X- and Y-directions, and 1.23 ± 0.16 µm in the Z-direction (median ± interquartile range, IQR), calculated over 4316 beads with diameter of 170 nm and imaged with a pixel size of 205 nm in sample space. Considering the angular overlap of the collection O1 and O2/3 collection cones [17], [27], the system has a latitudinal and longitudinal NA of 1.07 and 0.72 respectively, with O2/3 limiting the overall NA. For an emission wavelength of 525 nm, the estimated diffraction limited resolution of the microscope is 0.25 µm and 0.37 µm in the X and Y direction respectively. The median FWHM of the laterally-integrated axial intensity of the bead image was measured as 3.76 ± 0.79 µm. This is comparable to the light-sheet thickness of 3.5 µm (FWHM), measured using a Rhodamine 6G spin-coated fluorescent sheet and also to the calculated light-sheet FWHM of 3.5 µm, estimated from the initial beam diameter and system magnification.

**Figure 4.**
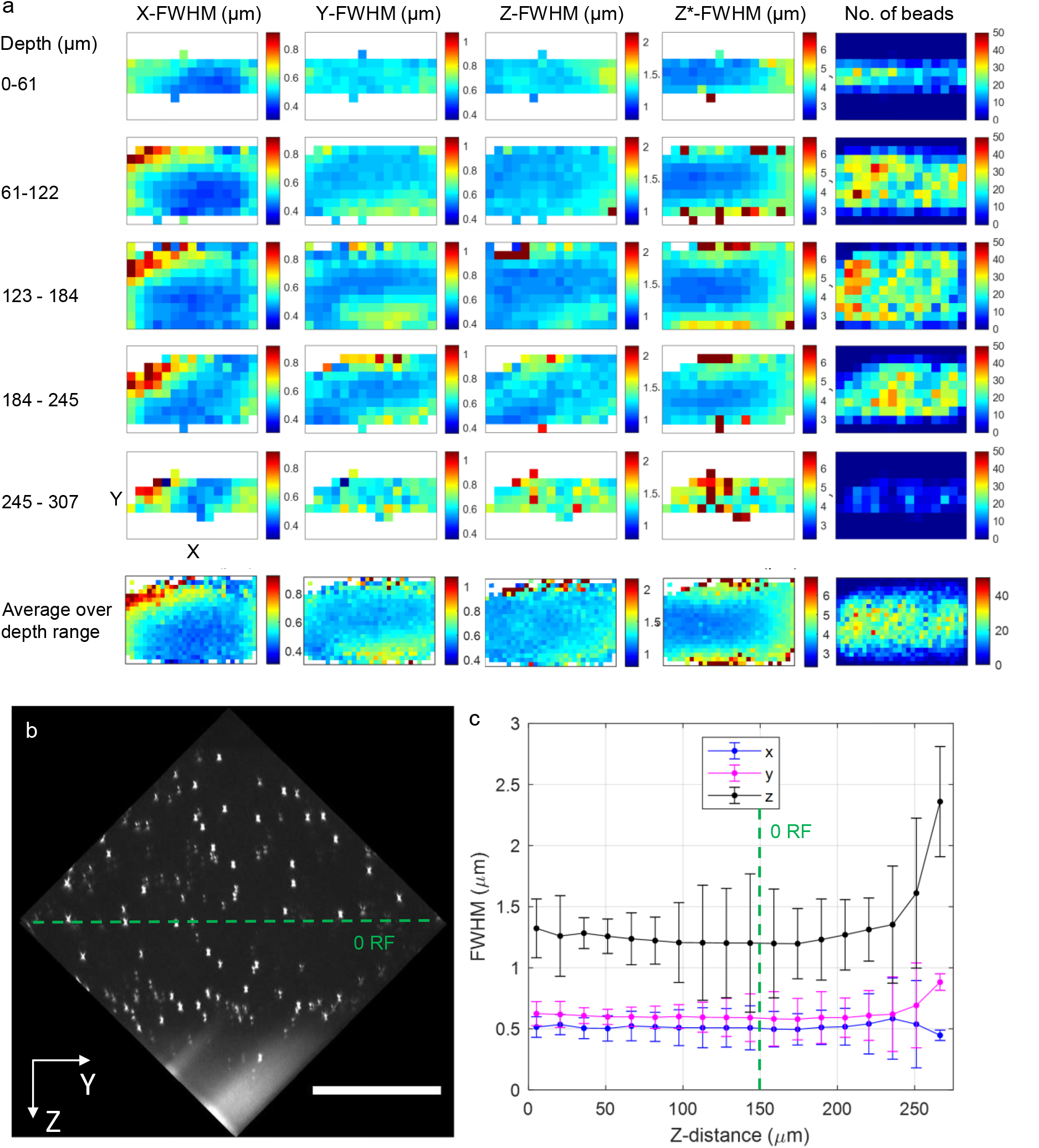
(a) Columns: Measured experimental lateral (X,Y) and axial (Z) PSF FWHM, optical sectioning strength (Z*-FWHM), and number of beads for a 480 µm × 320 µm XY-region, representing the median value for binned FWHM over 15 × 10 × 5 (X × Y × Z) voxel spatial regions with dimensions 32 µm × 32 µm × 60 µm. The bottom row shows the average over the whole depth, binned into 30 × 20 (X × Y) regions with dimensions of 16 µm × 16 µm. (b) YZ-orthoslice of the bead volume, cropped to only include the fused overlapping region of the two views. Scalebar: 100 µm. (c) Average raw bead image X-, Y- and Z-PSF FWHM (blue, magenta and black respectively) as a function of imaging distance from coverslip. Error bars represent the standard deviation. The dashed green line in (b) and (c) indicates the focal plane of O1, i.e. plane of zero remote refocus, 150 µm into the sample from the coverslip boundary.

#### 3.1.2. Correction collar adjustment

For each bead sample refractive index, we acquired a set of image volumes over a range of CC settings, see **Figure 5** and **Table S2**. For the sample with n = 1.4 (top row of **Figure *5***) we see a reversal in the spherical aberration sign with change in correction collar position about the nominal CC value of 0.17 (corresponding to 170 µm) as expected.

**Figure 5:**
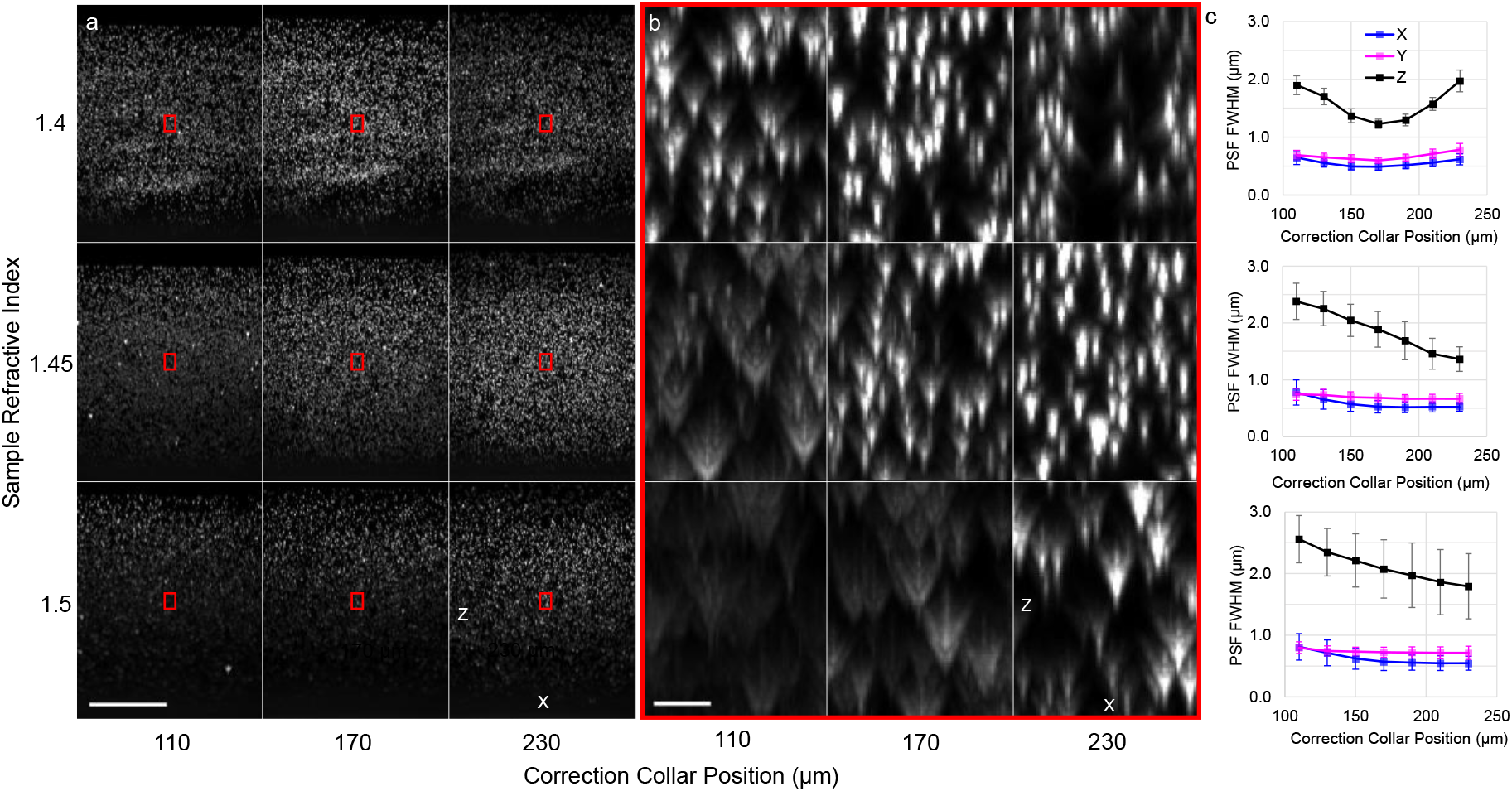
(a) X-Z MIP through fluorescent bead phantoms with a refractive index of 1.4 (top row), 1.45 (middle row) and 1.5 (bottom row). The columns represent the extreme and central value of the correction collar position of the primary objective. Scalebar: 100 µm. (b) Zoom-ins on the red regions in (a). Scalebar: 5 µm. (c) Measured experimental lateral (X,Y) and axial (Z) PSF FWHM for 1.4 (top), 1.45 (middle) and 1.5 (bottom) rows for varying correction collar positions, calculated as medians over the region of overlap between the two views. Error bars represent the inter-quartile range.

For the higher refractive index samples of 1.45 and 1.5, at the nominal CC value the images show depth-dependent spherical aberration (middle and bottom rows of Figure 5a), resulting in degraded axial resolution with increasing remote-refocus into the sample. We always see positive spherical aberration at the nominal CC value, as expected. While increasing the CC position can partially compensate for this effect, with less degraded Z-FWHM at higher CC values, the available CC range is insufficient to recover the axial resolution compared to the sample RI n = 1.4 (**Figure *5***c). The lateral resolution follows a similar trend, although the effect is less pronounced.

The change in lateral and axial resolution with imaging depth for the three refractive indices and three CC values is shown in **Figure S2**. For greater mismatch between the magnification of the remote refocusing relay and the sample RI, the axial resolution degrades faster with increased imaging depth. Again, we can see that adjustment of the CC position can partially compensate for the loss in axial resolution, but only over a limited axial range (∼50 µm).

### 3.2. Imaging thick biological samples of varying refractive indices

The system was used for proof-of-concept imaging of three different fixed biological samples (detailed preparation and mounting details in **Supplementary Notes 2-4**): mouse omentum containing metastatic murine ovarian carcinoma tissue slice (TDE-cleared and RI-matched to n = 1.406), a whole fly brain (uncleared, mounted in Vectashield (Vector Laboratories) with n = 1.45) and a mouse colon tissue slice (cleared and mounted in Ce3D, n = 1.5) (**Figure *6***). All animal experiments were carried out under approval of the local Ethical Review Committee according to UK Home Office guidelines. The correction collar was set to 0.17, 0.19 and 0.21 positions for the 1.406, 1.45 and 1.5 RI samples respectively, corresponding to partial compensation for the RI-mismatch to the design system magnification.

**Figure 6.**
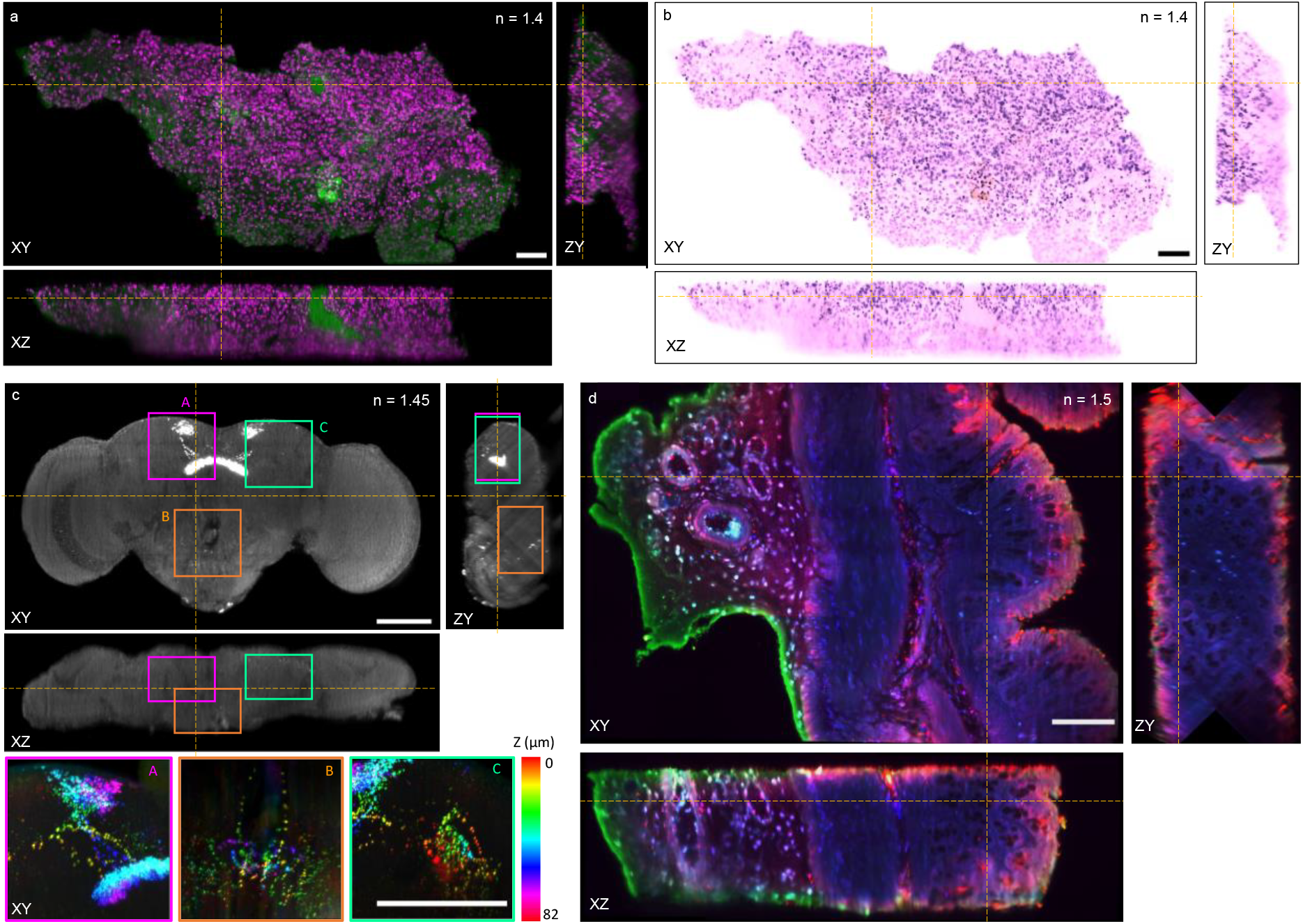
Orthogonal slices through tiled, stitched dual-view acquisitions of cleared biological samples with varying refractive indices, with the dashed yellow lines indicating the slice locations. (a) Mouse omentum tissue slice cleared and refractive-index matched with TDE, with DRAQ-7 labelled nuclei shown in magenta and autofluorescence shown in green. (b) False coloured version of the mouse omentum slice, imitating the H&E colour palette. c) *Drosophila melanogaster* fruit fly brain with *R23E10-GAL4-labelled* neurons expressing mitochondria-localized GFP. Gamma correction of 0.75 has been applied to aid visualization. The depth-encoded projections through three-dimensional regions of interest A (dendritic and axonal fields, in the superior protocerebrum and fan-shaped body), B (suboesophageal zone), C (cell bodies, in the posterior lateral protocerebrum) with dimensions of 122 µm × 122 µm × 82 µm (X×Y×Z) display the three-dimensional anatomical spread of neurons and the distribution of mitochondria therein. (d) Mouse colon tissue slice of 250 µm thickness, immunolabelled, and cleared using a modified Ce3D protocol [28]. Antibodies directed against E.coli gut bacteria are shown in green and blue, and the mucin marker Ulex Europaeus Agglutinin I is shown in red. Autofluorescence and/or non specific labelling are also evident across channels. All scalebars 100 µm.

The 1.6 mm × 0.8 mm, 0.25 mm thick mouse omentum tissue slice was imaged by stitching together an array of 8 × 4 dual-view fused tiles (**Figure *6***a). Each view consisted of 100 slices spaced by 2 µm, each slice acquired with a 15 ms exposure time, 2×2 pixel binning, with the dual-channel tiled acquisition totalling 17 minutes. Individual nuclei can be resolved across the whole tissue thickness. **Figure *6***b shows the same volume pseudo-coloured to mimic H&E-staining used in histopathology.

The *Drosophila melanogaster* fruit fly brain (**Figure *6***c) was acquired as a dual-view 3 × 2 tile acquisition with a 20 ms exposure time per slice, 2×2 pixel binning, and 121 slices per view and a z step of 2 µm. The FOV of the uncropped stitched volume covered 867 µm x 612 µm x 210 µm, enabling visualization of *R23E10-GAL4*^*+*^ neuronal distribution in 3D. The dual-view 6-tile acquisition covering the whole brain took 2 minutes 40 seconds to acquire.

The Ce3D-cleared colon tissue slice section (n = 1.5) (**Figure *6***d) was stitched from 3 × 2 tiles, each a fused dual-view acquisition. Various layers can be identified, with left to right: adipose tissue with blood vessels, muscle tissue, mucosa and the mucin barrier surrounding the lumen.

## 4. Discussion and Conclusions

In this work, we demonstrated the design and application of a dual-view oblique plane microscope for imaging optically cleared tissue. The remote refocusing relay was designed to operate with a silicone immersion primary objective, with the second tube lens composed of stock optics and aligned to achieve the target magnification invariant over a >400 µm defocus range as shown by imaging 3D distributions of fluorescent beads embedded in agarose. Using RI-tuned fluorescent bead samples and an automated 3D PSF characterization pipeline, the spatial resolution was quantified across the 3D FOV for n = {1.4, 1.45, 1.5} and the full range of possible collar correction positions. The system was then used for proof-of-concept imaging of thick tissue samples with the three different refractive indices, demonstrating fast high-resolution imaging over a large FOV.

While the remote refocusing relay design, together with CC adjustment enables imaging of samples with variable RIs, the image quality degrades with increasing RI mismatch to the design value. Zemax simulations with varied separations of the selected stock doublets that make up TL2 demonstrate that it is possible to achieve diffraction-limited performance over the TL2 focal length range needed to match sample refractive indices in the 1.33-1.55 range, however, such adjustment of TL2’s focal length also requires adjustment of other distances and overall path length of the remote refocusing relay. This problem has been addressed by Millet-Sikking [29] by replacing the static TL2 with a motorized dynamic zoom lens composed of three linearly translated lens elements enabling tuning of the magnification of the remote refocusing relay to the sample while maintaining a constant overall path length. The related approach of Prince et al. [30] involves the addition of an extra lens and mirror-based path compensator arrangement, with linear adjustment of the lens and mirrors positions allowing pre-calibrated tuning of the remote refocusing path length for compatibility with a selection of primary objectives. Nevertheless, our work shows that useful image resolution over a useful axial image range can be obtained by a simpler fixed TL2 arrangement over a range of sample refractive indices.

A further potential improvement would be motorizing the CC adjustment, as shown in Mohanan and Corbett [14], to compensate for the RI-mismatch of the sample to the design magnification of the remote refocusing relay, hence extending the usable refocusing range. This could be implemented automatically, using pre-calibrated collar positions based on a known sample refractive index and distance of the zero remote refocus plane beyond the coverslip.

Currently acquisitions of larger samples require tiling and stitching in both lateral directions. Implementing stage-scanning of the sample would enable unlimited FOV and no stitching artefacts in one of the lateral dimensions, reducing image processing time. These modifications would further extend the size of samples that the system can image while maintaining the sub-micron resolution.

In conclusion, we demonstrated a dual-view oblique plane microscope employing a silicone immersion primary objective and have characterised its performance when imaging bead samples with a range of refractive indices. The system achieves a lateral resolution of < 1 µm and an axial resolution of < 1.5 µm, maintained across >250 µmin imaging depth. We demonstrated that adjusting the collar correction on the primary objective can extend the usable imaging range for higher RI samples, making the system compatible with a range of clearing protocols. Our work presents a robust platform for fast volumetric imaging of few hundred µm thick optically cleared biological samples with subcellular resolution.

## 5. Author contributions

L.D., H.S., C.D. designed the system configuration. L.D. constructed and aligned the detection and illumination paths. R.S. prepared the fly brain sample. D.C. and D.T.R. prepared the cleared colon tissue sample. L.D., D.E., P.C., I.A.M. prepared the mouse omentum samples. L.D. acquired, processed and analyzed the data, drafted the manuscript and prepared the figures. All authors critically reviewed the manuscript.

## 6. Funding acknowledgments

Engineering and Physical Sciences Research Council (EPSRC, EP/T003103/1), Cancer Research UK (CRUK, A29386), and Ovarian Cancer Action (OCA, PSN417). L.D. was funded by a EPSRC Doctoral Prize Fellowship (EP/W524323). D.T.R. was funded by a Sir Henry Dale Fellowship (211230/Z/18/Z). P.C. is funded by a Parasol Foundation Fellowship.

