## Supplementary Information for "Cleared tissue dual-view oblique plane microscopy"

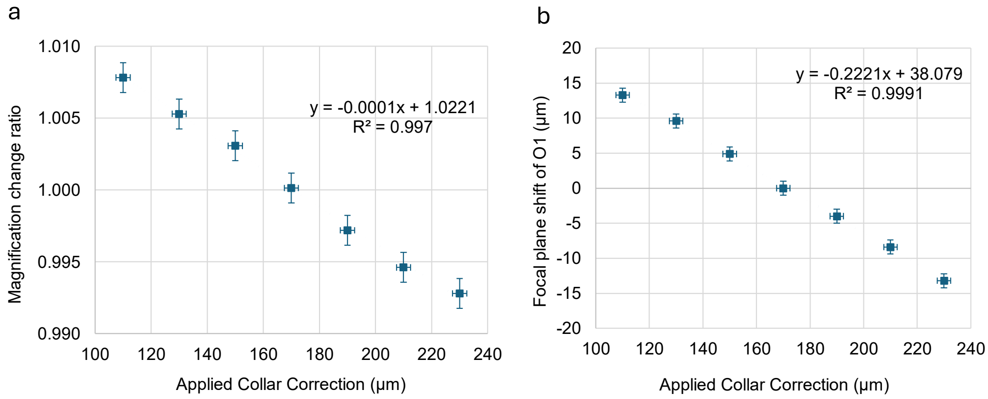

**Figure S1** (a) Change in the lateral magnification (average for x and y) of the first microscope (O1 and TL1) for different correction collar (CC) settings, with the available CC range causing a 1.5% change in the magnification. (b) Required axial translation of O1 to refocus on the sample after changing the correction collar value. Vertical error bars in (a) are the propagated errors based on single pixel dimensions, and in (b) correspond to the accuracy of in-focus readings of the motorized O1 z-drive position. Horizontal error bars correspond to the estimated errors of manual correction collar adjustment.

| **Sample** | **Bead phantoms** | **Mouse omentum** | **Fly brain** | **Mouse colon** |
|---|---|---|---|---|
| Clearing/mounting medium | TDE | TDE | Vectashield | Ce3D |
| RI | 1.406, 1.45, 1.5 | 1.406 | 1.45 | 1.5 |
| CC position | 0.11-0.23 | 0.17 | 0.19 | 0.21 |
| $Z_{ORF}$ (µm) | 150 | 100 | 100 | 150 |
| No of tiles (H × V) | 1 × 1 | 8 × 4 | 3 × 2 | 3 × 2 |
| Exposure time (ms) | 100 | 15 | 20 | 200 |
| Pixel binning | 1 × 1 | 2 × 2 | 2 × 2 | 1 × 1 |
| Z Step size (µm) | 1 | 2 | 2 | 1 |
| No of slices per view | 241 | 101 | 121 | 201 |
| Excitation channels (nm) | 488 | 488, 642 | 488 | 488, 561, 642 |

**Table S1** Summary of the image acquisition and processing parameters for each one of the samples for the data presented in the main text, including refractive index (RI), correction collar (CC) position, the distance of the plane of zero remote refocus to the coverslip surface, $Z_{ORF}$, number of horizontal (H) and vertical (V) tiles and their respective overlap, exposure time, pixel binning, Z step size, total number of slices per view, and laser excitation wavelength.

### Supplementary note 1: Preparation of RI-tuned fluorescent bead samples

The RI-tuned fluorescent bead phantoms were generated by mixing 2% agarose in liquid form with 2,2'-Thiodiethanol (TDE, Sigma Aldrich) to obtain the desired refractive index (1.406, 1.45 and 1.5), checked using a refractometer (Abbe 5 Refractometer, Bellingham and Stanley). Further agarose powder was added to allow the liquid to set into a gel when cooled, adjusting the final agarose concentration to ~2.2%, with the RI re-measured and adjusted through addition of TDE if necessary. Fluorescent beads with 170 nm actual diameter (Tetraspeck, Invitrogen T7280) were added to the agarose (heated to form a liquid) at 1:40 concentration, and after careful mixing, pipetted into a #1.5H glass bottom 8-well µ-slide (Ibidi, 80807).

### Supplementary note 2: Preparation of mouse ovarian cancer omentum samples

ID8-F3 mouse omental tumours were fixed in 10% neutral buffered formalin for at least 24 hours, then placed in 70% ethanol before embedding in 5% low gelling temperature agarose (Thermo Scientific) and macro-sectioned into 250 µm slices on a Leica VT1200S vibratome. The slices were washed and resuspended in phosphate-buffered saline (PBS) and labelled with the DNA stain DRAQ5 (Biostatus) at 10 µM concentration for 30 minutes. After further washes and resuspension in PBS, the samples were cleared and RI-matched through sequential incubation in TDE: 20% TDE/PBS for 2 hours in 37 degrees on a rocker, followed by 36.5% TDE/PBS (n=1.406) for 2 hours in 37 degrees on a rocker, and left on rocker at room temperature overnight. The samples were mounted in the second TDE dilution solution onto a microscope slide, using a gene frame to create a volume, and sealed with a #1.5 coverslip.

### Supplementary note 3: Preparation of fly brain samples

A detailed account of the fly brain sample preparation is provided in Sarnataro et al. (2025) [31]. Briefly, *Drosophila melanogaster* fruit flies expressing the mitochondria-localised fluorophore GFP (RRID: BDSC_8442) under the control of the driver line *R23E10-GAL4* (RRID: BDSC_49032) were reared on medium containing yeast, cornmeal, molasses, and agar under a 12 h light:12 h dark cycle at 25 °C with 60% relative humidity. Brains of randomly selected females were dissected with sharp forceps 4-6 days post eclosion, in phosphate buffer saline (PBS), quickly transferred to Eppendorf tubes on ice, and fixed in 4% (w/v) paraformaldehyde (Electron Microscopy Sciences) diluted in 0.3% (v/v) TritonX-100 in PBS (PBST) for 20 min on a nutator. Brains were quickly washed twice and then three times rocking for 20 min each, with PBST, and mounted in Vectashield antifade mounting medium (Vector Laboratories) on a microscopy slide (Fisherbrand), covered with a #1,5 coverslip (VWR), and sealed with nail polish.

### Supplementary note 4: Preparation of Ce3D-cleared colon tissue.

A frozen mouse colon sample was obtained from an experiment described in Carreno et al., 2024 [32]. This experiment was carried out under approval of the local Ethical Review Committee at Imperial College London according to UK Home Office guidelines (PPL PP7088487). Briefly, mice received an engineered commensal E. coli expressing constitutive fluorescent mKate2 and inducible mVenus to study gut inflammation. Dissected and fixed mouse colon was cryosectioned using a Leica Cryostat to remove the first 200 µm and expose the lumen for more efficient antibody permeability. Immunolabelling was performed using a primary Chicken IgY anti-GFP (Abcam) and Donkey anti-chicken IgY (H+L), CF™ 488A (Sigma-Aldrich) secondary antibodies for immunolabelling of mVenus; Goat IgG anti-RFP (Cambridge Biosciences) primary and Alexa Fluor 568 donkey anti-goat IgG (H+L) (Abcam) secondary antibodies for immunolabelling of mKate2, and Ulex Europaeus Agglutinin I (UEA I), DyLight® 649 (Vector Laboratories) preconjugated marker of the mucin layer. Ce3D clearing of mouse colon was adapted from Li et al., 2019 [28] as described in Carreno et al., 2024 [32]. The sample was placed in freshly made Ce3D clearing solution, protected from light in a cryomold for 10 days, replacing the Ce3D solution every 48h. Optically cleared mouse colon was mounted onto a microscope slide using gene frames to create a volume, Ce3D clearing solution as a mounting media (n = 1.5) and sealed with a glass cover slip with #1.5 thickness.

| Refractive index | Collar Correction (µm) | Number of Beads | X-FWHM (µm) | | Y-FWHM (µm) | | Z-FWHM (µm) | | Z*-FWHM (µm) | |
|---|---|---|---|---|---|---|---|---|---|---|
| | | | Median | IQR | Median | IQR | Median | IQR | Median | IQR |
| 1.4 | 110 | 4388 | 0.65 | 0.24 | 0.69 | 0.16 | 1.90 | 0.32 | 3.90 | 0.61 |
| | 130 | 4371 | 0.55 | 0.14 | 0.65 | 0.14 | 1.70 | 0.28 | 3.85 | 0.65 |
| | 150 | 4360 | 0.50 | 0.12 | 0.63 | 0.14 | 1.37 | 0.24 | 3.79 | 0.80 |
| | 170 | 4316 | 0.49 | 0.11 | 0.60 | 0.10 | 1.23 | 0.16 | 3.75 | 0.79 |
| | 190 | 4368 | 0.52 | 0.12 | 0.64 | 0.13 | 1.30 | 0.20 | 3.75 | 0.71 |
| | 210 | 4370 | 0.56 | 0.15 | 0.71 | 0.16 | 1.58 | 0.22 | 3.74 | 0.58 |
| | 230 | 4264 | 0.62 | 0.19 | 0.78 | 0.22 | 1.97 | 0.38 | 3.77 | 0.51 |
| 1.45 | 110 | 3324 | 0.78 | 0.44 | 0.74 | 0.21 | 2.38 | 0.64 | 4.15 | 0.51 |
| | 130 | 3961 | 0.65 | 0.35 | 0.73 | 0.19 | 2.25 | 0.61 | 4.02 | 0.52 |
| | 150 | 4454 | 0.57 | 0.26 | 0.69 | 0.19 | 2.05 | 0.56 | 3.91 | 0.56 |
| | 170 | 4447 | 0.52 | 0.21 | 0.68 | 0.17 | 1.89 | 0.63 | 3.82 | 0.65 |
| | 190 | 4637 | 0.51 | 0.19 | 0.67 | 0.15 | 1.69 | 0.67 | 3.76 | 0.73 |
| | 210 | 4642 | 0.52 | 0.18 | 0.66 | 0.16 | 1.46 | 0.54 | 3.72 | 0.76 |
| | 230 | 4558 | 0.52 | 0.15 | 0.67 | 0.19 | 1.36 | 0.44 | 3.73 | 0.79 |
| 1.5 | 110 | 894 | 0.81 | 0.43 | 0.80 | 0.19 | 2.56 | 0.77 | 4.36 | 0.58 |
| | 130 | 1074 | 0.72 | 0.42 | 0.74 | 0.16 | 2.35 | 0.77 | 4.24 | 0.46 |
| | 150 | 1426 | 0.62 | 0.34 | 0.73 | 0.15 | 2.21 | 0.86 | 4.11 | 0.48 |
| | 170 | 1726 | 0.57 | 0.28 | 0.73 | 0.16 | 2.08 | 0.95 | 4.02 | 0.51 |
| | 190 | 1992 | 0.56 | 0.24 | 0.72 | 0.18 | 1.97 | 1.05 | 3.94 | 0.57 |
| | 210 | 2304 | 0.55 | 0.24 | 0.72 | 0.20 | 1.86 | 1.06 | 3.87 | 0.67 |
| | 230 | 2528 | 0.55 | 0.23 | 0.71 | 0.21 | 1.79 | 1.06 | 3.81 | 0.69 |

**Table S2**: Measured experimental lateral (X,Y),axial (Z),PSF FWHM and optical sectioning strength (Z*) for 1.4 (top), 1.45 (middle) and 1.5 (bottom) rows for varying correction collar positions, calculated as medians and corresponding interquartile ranges for beads across the whole overlapping volume region.

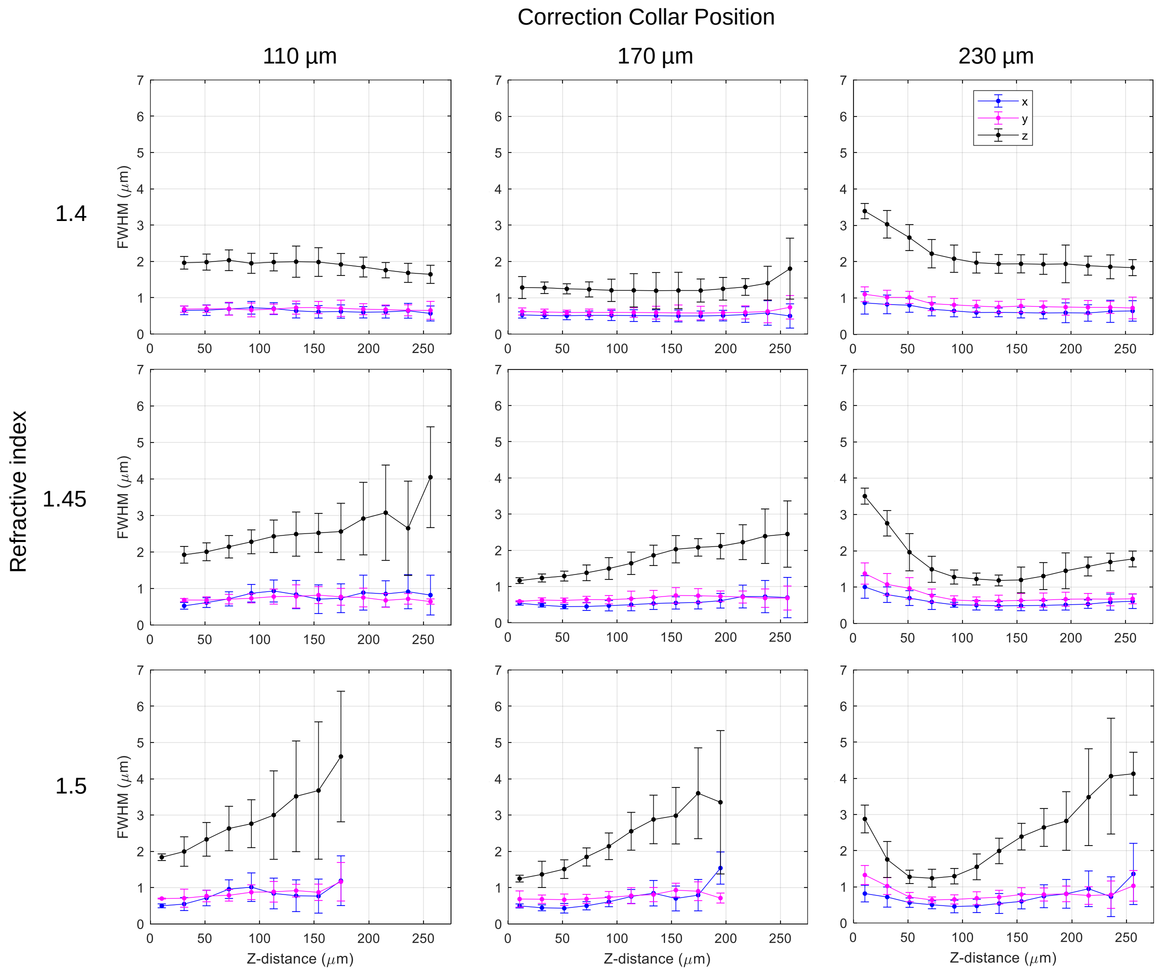

**Figure S2** Change in the average raw bead image X-, Y- and Z- PSF FWHM (blue, magenta and black respectively), with increasing imaging depth from the coverslip into the sample at various bead sample refractive indices (rows) and different correction collar positions (columns). Error bars represent the standard deviation. Varying offset of the initial data point for each series is due to shifted focus.
